# Neutrophils require SKAP2 for reactive oxygen species production following C-type lectin and *Candida* stimulation

**DOI:** 10.1101/2021.03.02.433609

**Authors:** Giang T. Nguyen, Shuying Xu, Stephen C. Bunnell, Michael K. Mansour, David B. Sykes, Joan Mecsas

## Abstract

Signaling cascades that convert the recognition of pathogens to efficient inflammatory responses by immune cells, specifically neutrophils, are critical for host survival. SKAP2, an adaptor protein, is required for reactive oxygen species (ROS) generation following stimulation by integrins, formyl peptide receptors and gram-negative bacteria *Klebsiella pneumoniae* and *Yersinia pseudotuberculosis in vitro* (Nguyen et al., 2020, Shaban et al., 2020, Boras et al., 2017). SKAP2 is also required for the host defense against *K. pneumoniae* and *ΔyopH Y. pseudotuberculosis* infection *in vivo* in mouse models (Shaban et al., 2020, Nguyen et al., 2020). Another class of pattern recognition receptors (PRR) is the C-type lectin receptors (CLR), such as Dectin-1, Dectin-2 and Mincle, that are critical to trigger innate immune responses. Using neutrophils from murine HoxB8-immortalized progenitors, we show that SKAP2 is crucial for maximal ROS response to purified CLR agonists and to the fungal pathogens *Candida glabrata* and *C. albicans*, as well as for robust killing of *C. glabrata*. *Skap2-/-* murine neutrophils failed to generate ROS and exhibited reduced cellular adhesion in response to trehalose-6,6’-dibehenate (TDB), furfurman, and curdlan, Mincle, Dectin-2, and Dectin-1 agonists, respectively. TDB, furfurman, and curdlan stimulation also led to SKAP2-independent integrin conformational changes, showing that inside-out signaling by these CLRs to integrin occurs in the absence of SKAP2. Pyk2 phosphorylation was significantly reduced after infection with *C. glabrata* in *Skap2-/-* neutrophils, while Syk phosphorylation was unaffected by the loss of SKAP2. These data strengthen the importance of SKAP2 in the activation of neutrophil ROS production by PRRs to include CLRs and extend the role of SKAP2 in host defense beyond antibacterial immunity to include *Candida* species.

## Introduction

Neutrophil inflammatory responses to infection are mediated by tyrosine kinase signaling cascades downstream of various pattern recognition receptors, including C-type lectin receptors (CLR), which are critical for host survival. The engagement of CLRs, Dectin-1, Dectin-2, or Mincle to bacterial and fungal pathogens, including *Klebsiella pneumoniae*, and *Candida* species, results in a diverse repertoire of antimicrobial functions in dendritic cells, macrophages, and neutrophils, including the generation of Dectin-1-mediated NADPH complex-derived reactive oxygen species (ROS) by neutrophils (Sharma et al., 2017, Sharma et al., 2014, Li et al., 2011, Wu et al., 2019, Thompson et al., 2019, Netea et al., 2015, Wells et al., 2008, Ifrim et al., 2014, Mansour et al., 2013, Hopke et al., 2020). Dectin-1 binds to β1,3-glucans, while Dectin-2 and Mincle bind to mannosylated ligands (Dambuza and Brown, 2015). CLR functions are regulated extensively by expression levels, localization, cooperation through heterodimerization, and crosstalk to β_2_ integrins (Ostrop and Lang, 2017, Lee et al., 2012, Li et al., 2011). Functional and biochemical analyses of CLRs in dendritic cells and macrophages have shown that their downstream signaling cascades involve the recruitment and phosphorylation of spleen tyrosine kinases (Syk) resulting in such antimicrobial functions including the release of ROS and pro-inflammatory cytokines to mediate innate and adaptive immune responses (Strasser et al., 2012, Ostrop et al., 2015, Rogers et al., 2005, Yamasaki et al., 2008, Saijo et al., 2010, Negoro et al., 2020).

Neutrophil ROS production is driven by signal transduction pathways downstream of several receptors, including integrin, Dectin-1, Fcγ, and G-protein-coupled receptors (GPCRs), that activate the components of the NADPH oxidase complex (Nguyen et al., 2017). Recently, we and others reported that <u>S</u>rc <u>K</u>inase <u>A</u>ssociated <u>P</u>hosphoprotein-2 (SKAP2) was essential for ROS production in response to stimulation by integrin and GPCRs, as well as gram-negative bacteria *K. pneumoniae* and *Yersinia pseudotuberculosis* (Nguyen et al., 2020, Shaban et al., 2020, Boras et al., 2017). In addition, SKAP2 is critical for integrin-stimulated cytoskeletal changes in macrophages and neutrophils (Alenghat et al., 2012, Tanaka et al., 2016, Boras et al., 2017), and for *K. pneumoniae*-mediated Syk phosphorylation in neutrophils (Nguyen et al., 2020). Further highlighting the role of SKAP2 in the innate immune response, *Skap2-/-* mice are more susceptible to *K. pneumoniae*, exhibiting a higher *K. pneumoniae* bacterial burden than wild-type controls (Nguyen et al., 2020) and SKAP2 is an essential target of the virulence factor, YopH, in neutrophils during *Y. pseudotuberculosis* infection (Shaban et al., 2020).

Here, we show that SKAP2 plays a critical role in pathogen recognition by another family of pattern-recognition receptors, the CLRs, and that Skap2 is critical for neutrophilic fungicidal response. Further, SKAP2, Src Family Kinases, and Btk are required for maximal ROS production following stimulation by *C. glabrata*, a fungal species that contributes to 27% of *Candida* bloodstream infections in the US, and is increasingly resistant to antifungal agents (Silva et al., 2012, Prevention, 2019). This study extends the importance of *Skap2* in host defense to additional microbial kingdoms and highlights the need to investigate the role of *Skap2* in human disease.

## Results and Discussion

### *Skap2* in neutrophils is dispensable for the expression of Mincle, Dectin-2, Dectin-1, and integrin receptors during pneumonic *K. pneumoniae* infection

To aid our studies of neutrophil functions, we used a granulocyte-monocyte progenitor (GMP) cell line that was conditionally immortalized by the enforced expression of an estrogen receptor-homeobox B8 (ER-Hoxb8) as previously described (Sykes and Kamps, 2001, Saul et al., 2019, Nguyen et al., 2020). The media for these cells contains stem cell factor and estradiol (E2), which permits nuclear translocation of ER-Hoxb8 fusion protein, resulting in the conditional maturation arrest at the GMP stage. The removal of E2 from the cell media, coupled with interleukin-3 (IL-3) and granulocyte colony stimulating factor (G-CSF) permits synchronous differentiation of GMP into mature neutrophils, which we termed differentiated *in vitro* (DIV) neutrophils (Nguyen et al., 2020). BALB/c wild-type and *Skap2-/-* DIV neutrophils were previously characterized and shown to be morphologically and functionally similar to their bone marrow (BM) counterparts (Nguyen et al., 2020).

To evaluate whether DIV neutrophils are functional *in vivo*, C57BL/6J (B6) wild-type and B6 *Skap2*−/− DIV neutrophils were transfused into neutropenic B6 mice, followed by intranasal infection with *K. pneumoniae* (ATCC 43816). Consistent with prior studies (Xiong et al., 2015, Nguyen et al., 2020, Ye et al., 2001), *K. pneumoniae* reached between 10^5^-10^6^ colony forming units (CFU) in wild-type (Figure 1A, control, black circles) and in neutropenic mice that received B6 wild-type DIV (Figure 1A, blue triangles) by 24 hours post-infection. However, mice that received B6 *Skap2-/-* DIV neutrophils (Figure 1A, red triangles) yielded a significantly higher 10^7^ CFU on average. The bacterial burden observed in neutropenic mice that received B6 DIV neutrophils (Figure 1A, blue triangles) was similar to what we previously observed in irradiated BALB/c mice that received wild-type BM cells that were depleted of inflammatory monocytes (Nguyen et al., 2020). This data indicates that *Skap2*-expressing DIV neutrophils are functionally comparable to BM-derived *Skap2*-expressing neutrophils *in vivo* in controlling bacterial burden, and that SKAP2-mediated neutrophil effector functions are critical for host protection against *K. pneumoniae* pneumonic infection.

**Figure 1:**
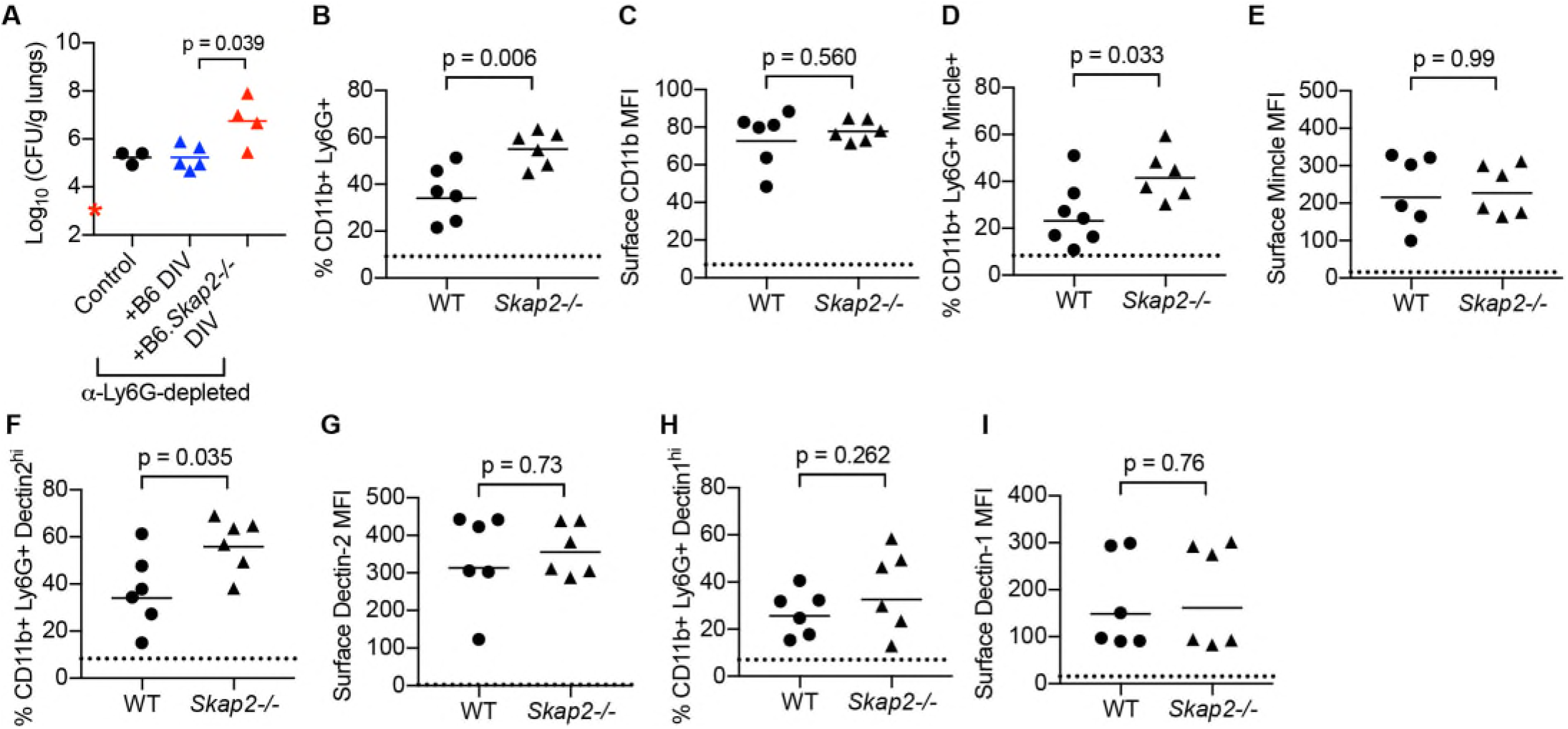
Transferred *Skap2-/-* DIV neutrophils fail to control *K. pneumoniae* infection in neutrophil-depleted mice despite normal C-type lectin receptor expression. **(A)** C57BL/6J (B6) wild-type mice were treated with PBS (control, black circles) or α-Ly6G (1A8, triangles) antibody intraperitoneally. 16 hours post-antibody injection, α-Ly6G-treated mice were intravenously injected with sterile PBS containing no cells (black triangles), B6 wild-type (blue triangles) and *Skap2-/-* (red triangles) DIV; DIV neutrophils were generated as described in the *Materials and Methods* and harvested at day 4 post-differentiation. All B6 mice were then infected intranasally with 5 × 10^3^ colony forming units (CFU) *pneumoniae.* Lungs were harvested after 24 hours and plated for bacterial burden (log (CFU/g lungs)). Data is compiled from 3 independent experiments with 1-3 mice each group. **(B-H)** BALB/c wild-type (WT) or *Skap2-/-* mice were infected intranasally with *K. pneumoniae,* and lungs were harvested after 24 hours. Cells were stained with CD11b and Ly6G, fixed with 4% formaldehyde, and stained with α-Dectin-1, α-Mincle, or α-Dectin-2 along with their respective secondary fluorescent-conjugated antibody (see *Materials and Methods*). Neutrophils (CD11b^+^ Ly6G^+^) were further gated to analyze expression of Dectin-1, Mincle, and Dectin-2. CD11b+ Ly6G+ cells were assessed for CLRs and CD11b integrin expression, and presented as **(B, D, F, H)** percent of cells, or **(C, E, G, I)** mean fluorescent intensity (MFI). Data is compiled from 2 independent experiments with 3 mice each group. Bars are geometric means, and each dot is a mouse. Dotted lines indicate **(B, D, F, H)** average level in PBS-treated lungs, or **(C, E, G, I)** MFI of sample stained with fluorescent-conjugated secondary antibody only. Significance was assessed using **(A)** one-way ANOVA with Sidak’s, or **(B-I)** two-tail unpaired Student’s *t* test.

Neutrophils from *K. pneumoniae*-infected lungs were previously shown to express Mincle receptors, which are critical for host defense and neutrophil function against *K. pneumoniae* (ATCC 43826) (Sharma et al., 2017, Sharma et al., 2014). To determine if other C-type lectin receptors are potentially playing a role in the pathogenesis of *K. pneumoniae* pneumonia, we surveyed the expression of Dectin-2 and Dectin-1 in the lungs of mice by flow cytometry (Figure S1). Consistent with prior data (Sharma et al., 2014), 99% of the highly Mincle-positive cells in the lungs of *K. pneumoniae* infected mice were CD11b^+^ Ly6G^+^ neutrophils (Figure S1A). In addition, flow cytometry showed lung cells expressing variable levels of Dectin 2 and Dectin-1, with the majority of cells that expressed high levels of Dectin-1 and Dectin-2 being CD11b^+^ Ly6G^+^ neutrophils (Figure S1B-C). These data show that Mincle, Dectin-1, and Dectin-2 are expressed on neutrophils, and may play a role in the pathogenesis of *K. pneumoniae* pneumonia. We next evaluated if the increased susceptibility of *Skap2-/-* mice to *K. pneumoniae* infection (Figure S1D) was due to differences in the expression of CLRs on neutrophils in *K. pneumoniae*-infected BALB/c wild-type (WT) and *Skap2-/-* lungs. Although there were more neutrophils recovered from *K. pneumoniae-*infected *Skap2-/-* lungs than WT lungs, there was no difference in the surface expression of CD11b (Figure 1B-C, Figure S1E), consistent with our prior DIV neutrophil data (Nguyen et al., 2020). Higher numbers of Mincle- and Dectin-2-positive cells and equivalent numbers of Dectin-1 expressing cells were recovered from *K. pneumoniae*-infected *Skap2-/-* lungs (Figure 1D-I, Figure S1F-H). Collectively, these data shows the CLRs are present on surfaces of neutrophils during *K. pneumoniae* infection independent of the presence of SKAP2, so the increased bacterial burden observed in *Skap2-/-* mice was likely not due to the lack of these receptors.

### SKAP2 modulates the activation of ROS production downstream of TDB, furfurman, and curdlan stimulation

ROS is required for host protection against *K. pneumoniae.* Patients with Chronic Granulomatous Disease generate less ROS than healthy hosts, and are highly susceptible to life-threatening *K. pneumoniae* respiratory infections (Bortoletto et al., 2015, Wolach et al., 2017), while ROS defective mice (B6-*Cybb−/−)* suffer from higher bacterial burdens than B6 mice (Nguyen et al., 2020, Paczosa et al., 2020). We examined whether *Skap2-/-* neutrophils are defective for ROS production following stimulation with the purified ligands trehalose-6,6’-dibehenate (TDB), furfurman, or curdlan, which stimulate Mincle homo/heterodimers, Dectin-2 and Dectin-1 homodimers, respectively. A cytochrome C absorbance assay was used to detect superoxide production (Nguyen et al., 2020, Dahlgren et al., 2020, Shaban et al., 2020). DIV and BM BALB/c wild-type (WT) neutrophils stimulated with TDB, furfurman, or curdlan produced significantly higher levels of superoxide compared with *Skap2-/-* counterparts (Figure 2A-B, Figure S2). The loss in ROS production was not due to defects in NADPH oxidase as *Skap2-/-* BM and DIV neutrophils robustly released ROS following PMA stimulation (Figure 2A), or to changes in the expression of Mincle, Dectin-2, and Dectin-1 on the surface of *Skap2-/-* DIV neutrophils (Figure 2C-H).

**Figure 2:**
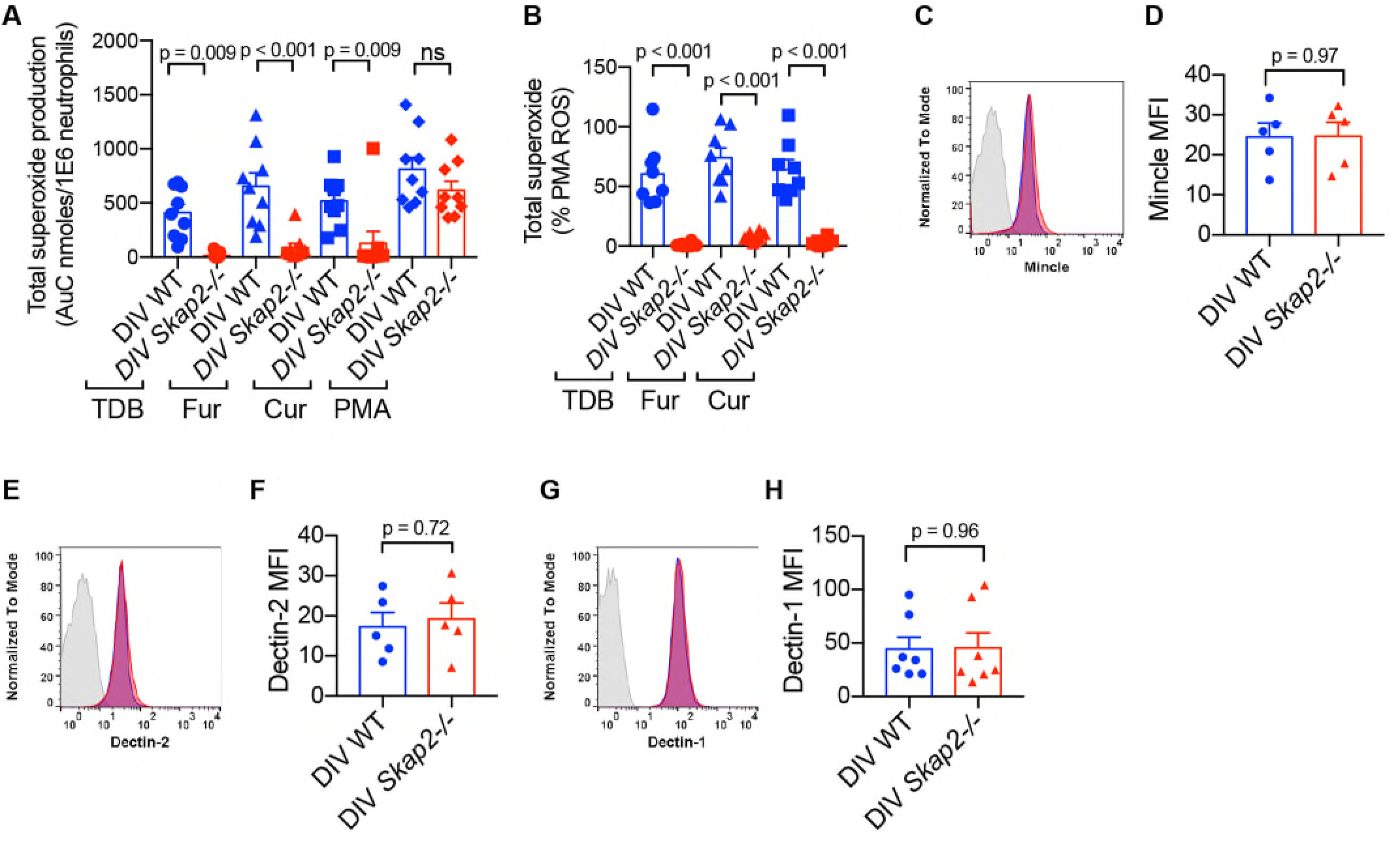
TDB, furfurman, and curdlan-induced ROS requires SKAP2. **(A-B)** The respiratory burst of BALB/c wild-type (WT) and *Skap2-/-* DIV neutrophils plated on immobilized TDB, an agonist for mincle-containing dimers, furfurman (Fur), an agonist for Dectin-2, or curdlan (Cur), an agonist to Dectin-1. The total concentration of superoxide produced after 60 minutes was calculated by the sum of the area under the curves, and presented as **(A)** total superoxide (values were normalized to unstimulated samples of each respective genotype), or **(B)** as percentage of PMA-induced ROS. Data are presented as mean + SEM compiled from 8-9 independent experiments from two different HoxB8-GMP cell lines per each genetic background (WT or *Skap2*−/−). **(C-H)** DIV neutrophils were stained with CD11b and Ly6G, fixed with 4% formaldehyde, and stained with α-Mincle, α-Dectin-2, or α-Dectin-1. Surface receptor levels are reported as mean fluorescence intensity (MFI) with mean + SEM from 5-7 independent experiments from two different HoxB8-GMP cell lines. Significance was assessed using **(A-B)** one-way ANOVA with Sidak’s, or **(D, F, H)** two-tail unpaired Student’s *t* test.

### SKAP2 is required for TDB, furfurman, and curdlan-stimulated cell adhesion

CLR activation triggers cell adhesion, spreading, and actin polymerization (Lee et al., 2012, Lee et al., 2017). These processes may influence ROS production, as the regulatory components of the NADPH oxidase complex interact with the actin cytoskeleton in both resting and activated neutrophils (el Benna et al., 1994, Nauseef et al., 1991, Woodman et al., 1991). Therefore, we examined whether SKAP2 is required for neutrophil spreading following TDB, furfurman, and curdlan stimulation. After one hour of TDB, furfurman, or curdlan stimulation, WT neutrophils adhered strongly to agonist-coated surfaces with 40-60% of the cells spreading to the agonist-coated wells in contrast to cells plated onto unstimulated surfaces (Figure 3A-D, K-L). However, the percentage of *Skap2-/-* neutrophils that spread on CLR agonist-coated surfaces was 3 to 4-fold less (Figure 3F-I, K-L). The reduction in cell spreading was not due to global defects in cytoskeletal rearrangement as *Skap2-/-* DIV neutrophils were able to spread on surfaces coated with IgG immune complexes (IC) albeit at only 60% of WT levels (Figure 3E, J-L). The reduced IC-induced spreading observed in *Skap2-/-* neutrophils was consistent with a previously reported reduction in IC-stimulated ROS production (Nguyen et al., 2020, Shaban et al., 2020). These data indicate that SKAP2 plays a role in CLR-induced cytoskeletal rearrangement leading to adhesion, and that the reduction in SKAP2-dependent, CLR-stimulated cytoskeletal remodeling in *Skap2-/-* neutrophils may hinder ROS production.

**Figure 3:**
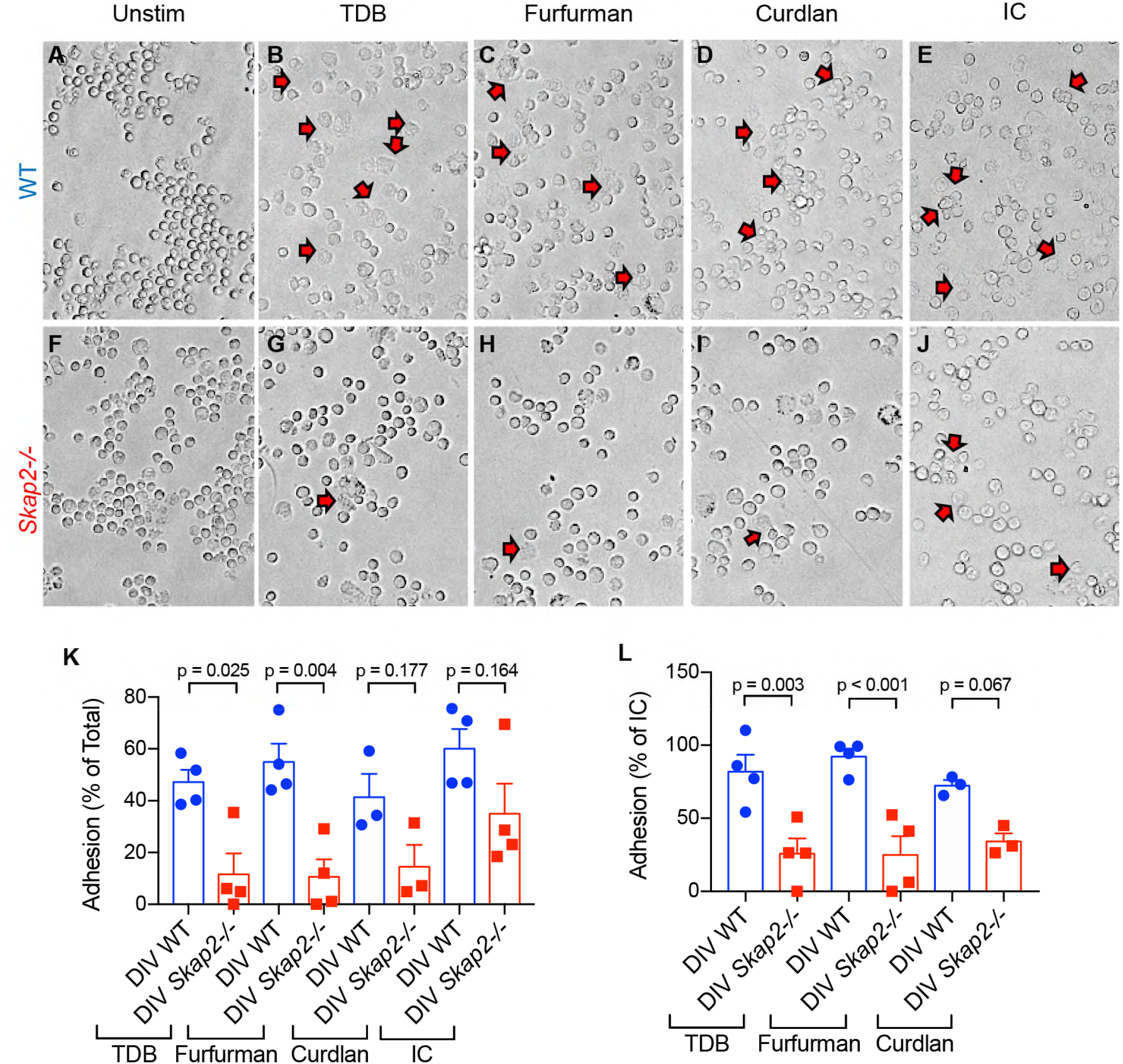
TDB, furfurman, and curdlan-stimulated neutrophil adhesion are SKAP2-dependent. **(A-J)** Firmly-adherent WT **(A-E)** and *Skap2-/-* **(F-J)** DIV neutrophils stimulated with immobilized TDB **(B, G)**, furfurman **(C-H)**, or curdlan **(D, I)** for 1 hour were imaged by bright-field microscopy. Control cells were plated onto surface coated with 10% FBS (Unstim) **(A, F)**, or IgG immune complexes (IC) **(E, J)** for 1 hour. Original magnification was 20x (n=3-4 in technical duplicates). **(K-L)** Images were blinded prior to counting the number of rounded and spread cells, which are presented as the percentage of **(K)** total cells counted (values were normalized to unstimulated samples of each respective genotype), or **(L)** of IC-induced adhered neutrophils. Significance was assessed using two-way ANOVA with Sidak’s post-test.

### SKAP2 is not required for TDB, furfurman, or curdlan-stimulated integrin activation

In neutrophils, Dectin-1 activates integrin receptors in response to zymosan, another Dectin-1 agonist, causing integrins to undergo conformational changes from inactive to high affinity states (Li et al., 2011). High affinity integrin states trigger ROS production, and can be assessed by rapid and inducible binding to ligands such as fibrinogen, which binds to β_2_ (CD11b/CD18) integrins (Van Strijp et al., 1993, Boras et al., 2017, Fagerholm et al., 2019). To determine if Mincle and Dectin-2 stimulation also induced an integrin conformational change to the high affinity form in neutrophils, a flow cytometric binding assay with Alexa-488-conjugated fibrinogen was used (Figure 4A-F). About 60-80% of murine neutrophils bound fibrinogen following treatment with Mn^2+^, the positive control for integrin activation (Figure 4C) (Dransfield et al., 1992, Ye et al., 2012). TDB, furfurman, or curdlan treatment of WT neutrophils resulted in integrin activation at approximately 40%, 30%, and 30% of Mn^2+^ level, respectively (Figure 4D-G). To determine whether SKAP2 is required to induce integrin conformational changes, we assessed whether *Skap2-/-* neutrophils had defects in CLR-induced integrin activation. We observed that similar percentages of *Skap2-/-* DIV neutrophils bound to Alexa-488-conjugated fibrinogen following TDB, furfurman, or curdlan stimulation indicating that CLR-induced integrin conformational changes are not SKAP2-dependent (Figure 4D-H). Combined, these data suggest that SKAP2-dependent defects in ROS are due to defects in the signaling cascades downstream of CLRs or integrins rather than changes in integrin activation.

**Figure 4:**
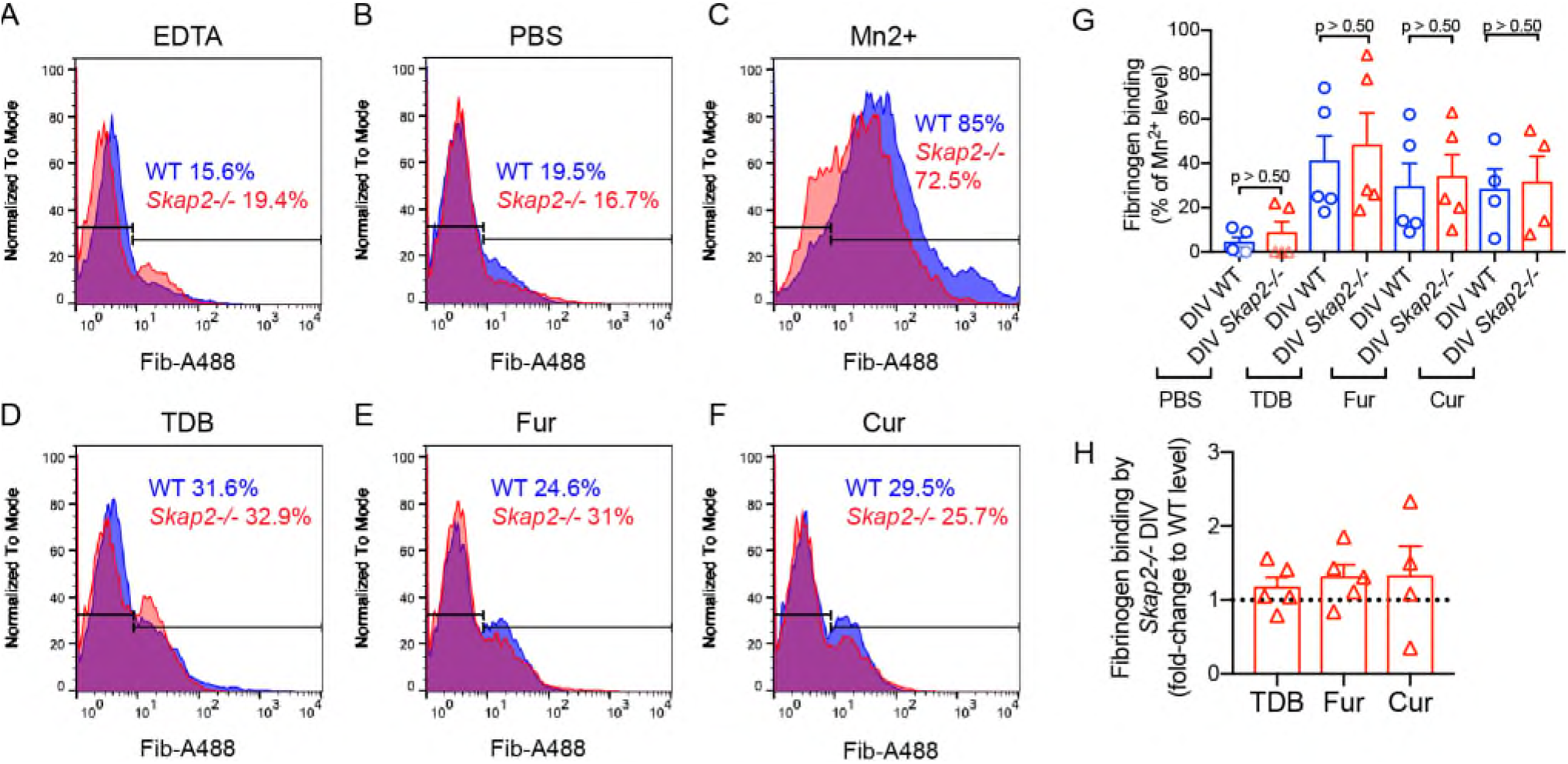
SKAP2 is not required for CLR-induced integrin activation. The assessment of high affinity integrin conformation was measured by the binding of soluble Alexa-488-conjugated fibrinogen to WT and *Skap2-/-* neutrophils stimulated with TDB, furfurman (Fur), and curdlan (Cur). Neutrophils were stimulated for 20 minutes at 37°C with 5% CO_2_. Alexa-488-conjugated fibrinogen was then added to each well, and the cells were incubated for another 15 minutes at 37°C with 5% CO_2_. Some cells were left untreated/unstimulated (PBS), treated with 5mM Mn^2+^ as a positive control for the last 15 minutes of incubation, or treated with 2.5mM EDTA for 30 minutes prior to the addition of fibrinogen to the plate as negative control. Cell suspensions were collected, fixed with 1% formaldehyde, and analyzed by flow cytometry. **(A-F)** Representative histograms gated on live cells. **(G)** The quantification of the percentage of fibrinogen-bound cells by DIV neutrophils (normalized to EDTA-treated level, divided by % of fibrinogen-bound by Mn2+-treated cells). **(H)** Level of fibrinogen-bound cells by *Skap2-/-* DIV neutrophils as fold-change to WT level. The data is presented as the mean + SEM from 4-5 independent experiments in technical duplicate. Significance was assessed using two-way ANOVA with Sidak’s post-test.

### SKAP2 is required for optimal *C. glabrata* and *C. albicans-* stimulated ROS production and overall fungal killing

Dectin-1, Dectin-2, Mincle, and β2 integrin receptors have all been individually proposed to be critical for fungal resistance in mice and humans (Chen et al., 2017, Feinberg et al., 2017, Ifrim et al., 2014, Sato et al., 2006, Wu et al., 2019, Saijo et al., 2010, McGreal et al., 2006, Zhu et al., 2013, Brown et al., 2003, Taylor et al., 2007, Ferwerda et al., 2009). The leading cause of invasive fungal infections involves *Candida* species including the morphologically and genetically different species, *C. albicans* and *C. glabrata,* that together account for 50-90% of cases in North America and Europe with a 40-50% mortality rate (Arendrup et al., 2011, Tortorano et al., 2006, Prevention, 2019). These three CLRs contribute to *C. albicans*-induced neutrophil ROS production, and are critical for the clearance of *C. albicans* (Thompson et al., 2019). Dectin-1 engagement leads to the activation of integrin receptors, resulting in neutrophil ROS production and protection against systemic *C. albicans* infection in mice (Li et al., 2011).

To dissect the contribution of SKAP2 in *C. glabrata* and *C. albicans*-activated neutrophils, the ROS production by *Skap2-/-* DIV neutrophils was assessed using isoluminol chemiluminescence (Nguyen et al., 2020, Dahlgren et al., 2020, Shaban et al., 2020). When stimulated with *C. glabrata* and *C. albicans*, WT DIVs generated robust and persistent ROS while *Skap2*−/− DIVs generated 50% and 25% of the levels produced by WT DIVs, respectively (Figure 5A-C). Next, we examined whether the defects in ROS affected the fungal killing ability by *Skap2-/-* DIV neutrophils. DIV neutrophils were incubated with *C. glabrata* for 24 hours, or *C. albicans* for 2 hours, and then fungal survival was measured using the PrestoBlue viability dye conversion (Xu et al., 2018, Tam et al., 2019). Compared to WT, *Skap2-/-* DIV neutrophils demonstrated significantly decreased fungicidal activity after infection with *C. glabrata* (Figure 5D-E) while infection with *C. albicans* typically also led to a decrease in fungal killing in 4 out of 5 experiments (Figure 5D, F). Combined, these data indicate that the SKAP2 deficiency resulted in an abrogated neutrophil ROS production after infection with *C. glabrata* and *C. albicans* that potentially enables fungal escape with *C. glabrata* appearing more sensitive to SKAP2-mediated responses.

**Figure 5:**
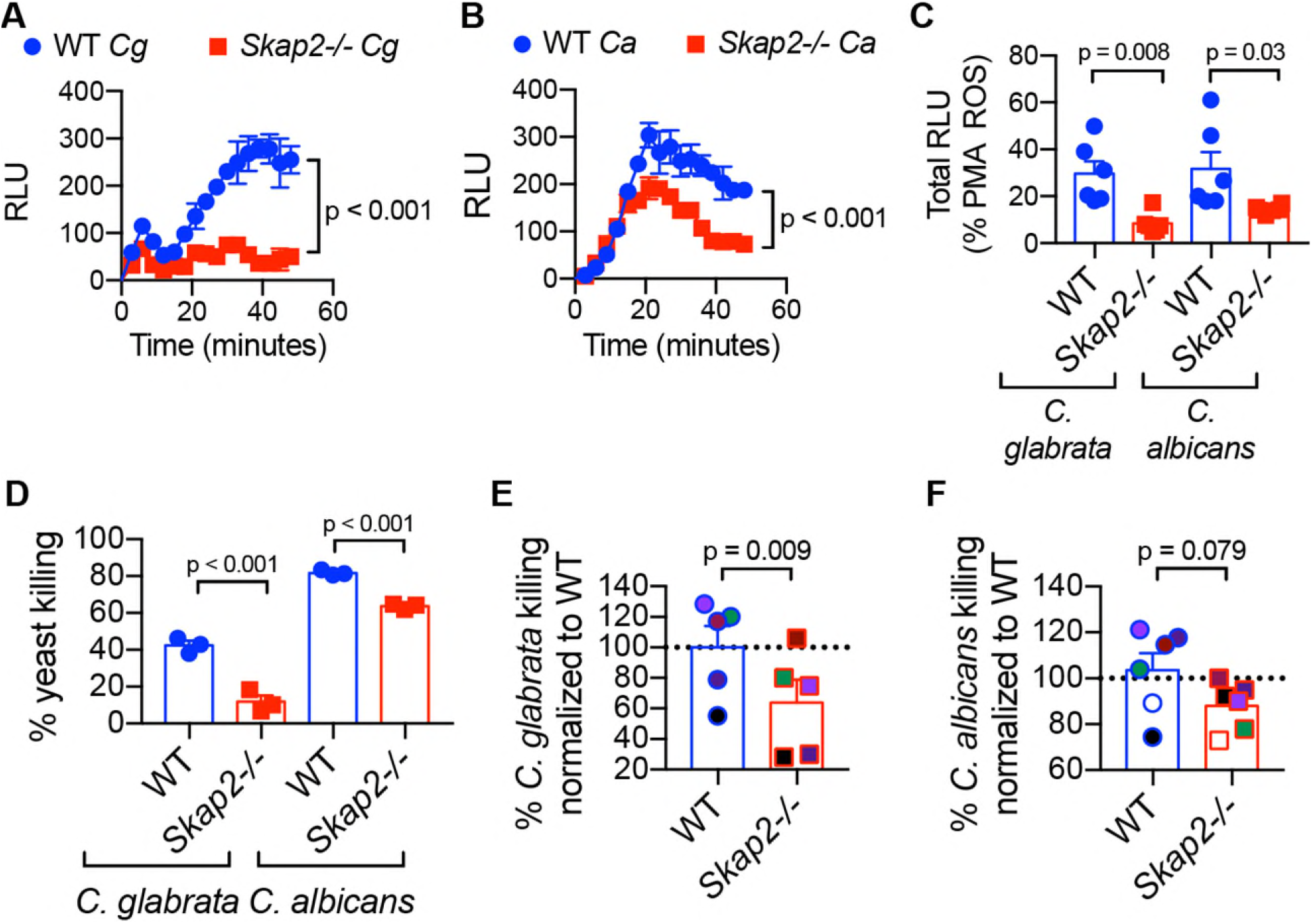
SKAP2 is required for maximal *C. glabrata* and *C. albicans*-induced ROS production, and neutrophil killing. **(A-C)** The respiratory burst of WT and *Skap2-/-* DIV neutrophils following *C. glabrata* and *C. albicans* stimulation (MOI 1) using isoluminol chemiluminescence. **(A-B)** A representative experiment performed in technical triplicate. Values shown are with background subtraction of the unstimulated samples plated on FBS-coated surface. **(C)** The total concentration of superoxide produced after 45 minutes was calculated by the sum of the area under the curves and presented as the percentage of PMA-induced ROS. DIV data are presented as mean + SEM compiled from n=6. Significance was assessed using **(A-B)** two-way ANOVA with Tukey’s post-test **(C)** one-way ANOVA with Sidak’s. **(D-F)** WT and *Skap2-/-* DIV neutrophils were co-incubated with yeasts (*C. albicans* for 2 hours, and *C. glabrata* for 18 hours) at MOI 0.2. Neutrophils were lysed and PrestoBlue dye was then added to measure the yeast viability by fluorescent readings. Percent of yeast killing was calculated as follows: 100 – (100*viability of yeast plus neutrophils /viability of yeast only). **(D)** Data is a representative in technical triplicate, and significance was assessed using two-way ANOVA with Sidak’s. **(E-F)** The average yeast killing of technical replicates was calculated from each independent experiment. The *Skap2-/-* values were divided by WT values in each respective experiment. The WT level of yeast killing in each experiment was set to 100 as indicated by the dotted line. The percent of yeast killing by WT DIV neutrophils was divided by the average of the 5-6 experiments. Data is compiled from the 5-6 independent experiments and presented as mean + SEM with each experiment represented by a different color. Significance was assessed using paired Students t-test.

### SKAP2 is required for optimal *C. glabrata*-stimulated phosphorylation of Pyk2, but not Syk kinases

Previous studies demonstrate Syk involvement in neutrophil ROS production in response to stimulation by Dectin-1 agonists, *C. albicans,* and *C. glabrata* (Deng et al., 2015, Negoro et al., 2020, Li et al., 2011). Consistent with these data, pretreatment of WT DIV with a small molecule inhibitor of Syk, R406, produced significantly lower TDB-stimulated ROS than their vehicle-treated counterpart (Figure 6A-B). Likewise, pretreatment of human neutrophils with R406 inhibited TDB- and Furfurman-stimulated ROS production (Figure 6C-D). Inhibitors of Src Family Kinases (SFKs), which act upstream of Syk, and of Bruton’s Tyrosine Kinase (Btk), which mediates phospholipase C activation downstream of Syk, also inhibited TDB-, Furfurman and curdlan-induced ROS production (Figure 6A-D) consistent with prior data in human neutrophils using PP2 (iSFK) and β-glucan beads (Nani et al., 2015). While Btk has been shown to contribute to neutrophil ROS, phagocytosis, and overall mouse survival during *C. albicans* infection (Colado et al., 2020, Strijbis et al., 2013), the role of SFKs and Btk in *C. glabrata* infection is unclear. Here, inhibitors of Syk, SFKs, and Btk also significantly reduced *C. glabrata-* and *C. albicans*-induced ROS production by DIV neutrophils (Figure 6E-F), suggesting that in addition to Syk (Negoro et al., 2020) and SKAP2 (Figure 5), *C. glabrata*-stimulated ROS also requires SFKs and Btk.

**Figure 6:**
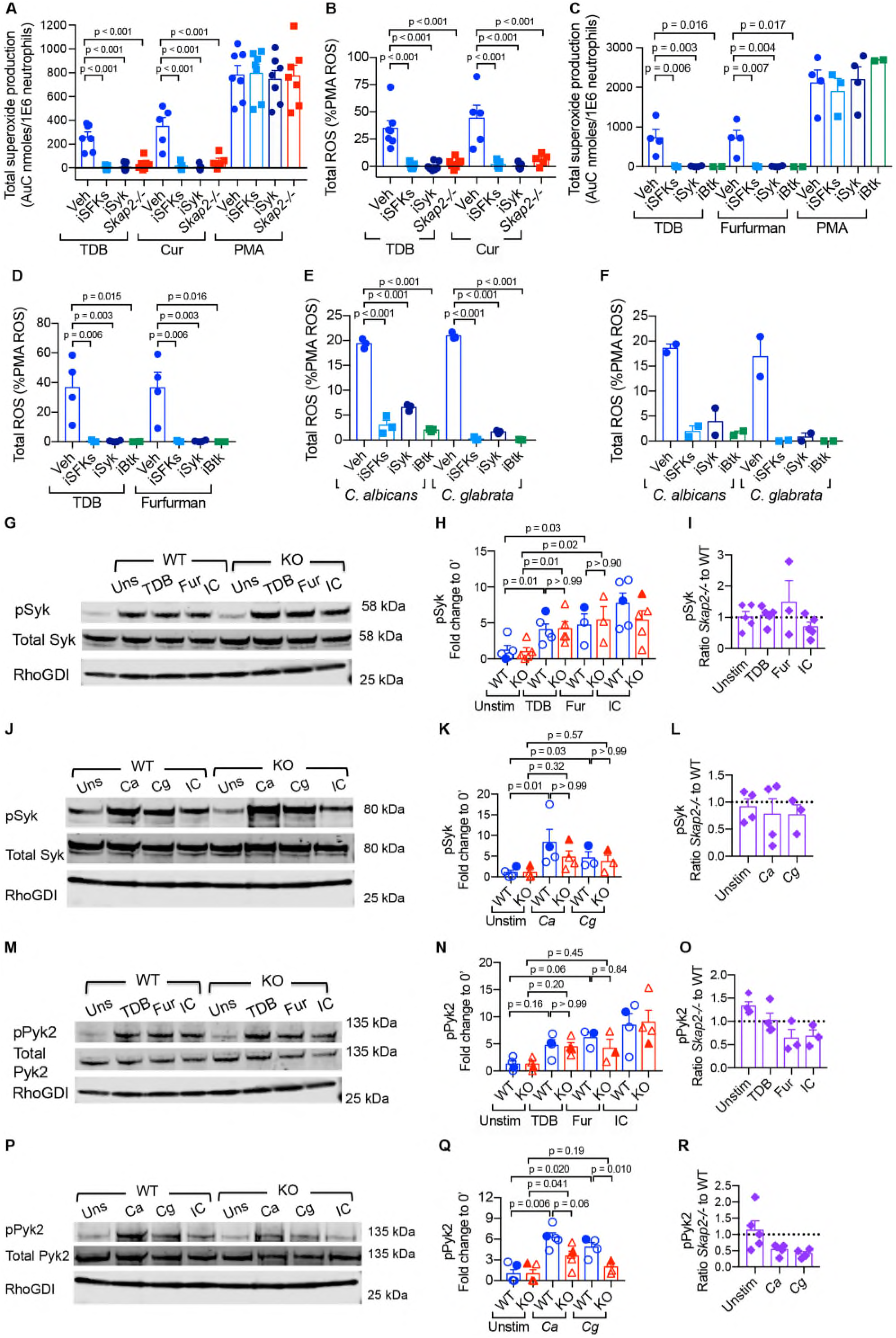
SFK-Syk-Btk are required for neutrophil ROS production following stimulation of TDB, curdlan, furfurman, *C. albicans*, and *C. glabrata*. The respiratory burst of DIV **(A-B, E-F)** or human **(C-D)** neutrophils treated with inhibitors using **(A-D)** a cytochrome C assay or **(E-F)** isoluminol chemiluminescence. **(A-D)** Neutrophils were pre-treated with DMSO (Veh), PP2 (iSFKs), R406 (iSyk), or Ibrutinib (iBtk) for 10 minutes at 37°C and then plated onto immobilized TDB, furfurman, or curdlan (cur), and measured for 60 minutes. The total superoxide was calculated as area under the curve after subtracting the uninfected samples, and presented as **(A, C)** total production or **(B, D-F)** as percentage of PMA-induced ROS. **(E-F)** DIV neutrophils were pre-treated with inhibitors as in (**A-D)**, and then infected with *C. albicans* or *C. glabrata* (MOI 1) and measured for 30 minutes. Total ROS production is presented as percentage of PMA-induced ROS. **(E)** A representative experiment is shown in technical triplicate. **(A-D, F)** Data is compiled from **(B-D)** 4-7 or **(F)** 2 independent experiments performed in technical triplicate and presented as mean + SEM with dots representing averages of independent experiments. **(A-F)** Significance was assessed using one-way ANOVA with Sidak’s post-test. **(G-R)** WT and *Skap2-/-* neutrophils were **(G-I, M-O)** plated onto immobilized TDB, furfurman (Fur), or IgG immune complexes (IC), or **(J-L, P-R)** stimulated with MOI 2 of *C. albicans* and *C. glabrata* for 15 minutes at 37°C. Lysates were analyzed by western blot for pSyk (Y352), pPyk2 (Y402), and RhoGDI. Blots were then stripped and re-probed for total Syk, and Pyk2. Phospho-signal was first divided by total protein, then by loading control, and then normalized to unstimulated values as described in the *Materials and Methods* and Supplementary file 1. Data is compiled from 3-5 independent experiments. **(G, J, M, P)** Representative blot shown. **(H, K, N, Q)** Fold change from each experiment with respect to unstimulated, symbols represent the value from each experiment, solid symbols indicating values of blot shown, bars indicate mean. Statistics represent mean + SEM. **(I, L, O, R)** The *Skap2-/-* values were divided by WT values in each respective experiment. The average WT level of CLR-stimulated phosphorylation was set to 1, as indicated by the dotted line. Significance was assessed using two-way ANOVA with **(B, E, H, K)** Tukey’s post-test or **(N, Q, T, W)** Sidak’s post-test

Previously, we showed that SKAP2 contributes to *K. pneumoniae*-induced activation of tyrosine kinases Syk and Pyk2 (Nguyen et al., 2020), which also function downstream of integrin receptors (Mocsai, 2002). To determine whether SKAP2 was required for the phosphorylation of Syk and Pyk2 following TDB, furfurman, *C. glabrata* or *C. albicans* stimulation, western blots of lysates from TDB, furfurman, or *Candida*-challenged WT and *Skap2-/-* DIV neutrophils were analyzed. Consistent with prior work in BM neutrophils using zymosan, a Dectin-1 agonist (Li et al., 2011), WT DIV neutrophils stimulated with TDB, furfurman, *C. albicans*, and *C. glabrata* enhanced the phosphorylation of Syk and Pyk2 relative to unstimulated controls (Figure 6G-R, Figure S3A-C, Table S1-4). However, in contrast to infection with *K. pneumoniae* (Nguyen et al., 2020), the loss of SKAP2 deficiency did not significantly reduce Syk phosphorylation downstream of TDB, furfurman, *C. glabrata* and *C. albicans* stimulation (Figure 6G-L, Figure S3A-C, Table S1, S2), or TDB and furfurman-induced Pyk2 phosphorylation (Figure 6M-O, Figure S3C Table S3-S4). On the other hand, *C. glabrata* and *C. albicans*-induced Pyk2 phosphorylation was reduced in *Skap2-/-* DIV neutrophils with the SKAP2-defect more pronounced in *C. glabrata*-stimulated conditions (Figure 6P-R, Figure S3A-C, Table S2, S4). In summary, after infection with *C. glabrata* and *C. albicans*, Pyk2, but not Syk phosphorylation is reduced in absence of SKAP2 whereas after infection with *K. pneumoniae,* the phosphorylation of both kinases requires SKAP2 (Figures 6, Figure S3C) (Nguyen et al., 2020).

Collectively, our results show that SKAP2 plays a critical role in optimal neutrophilic fungicidal responses to *C. glabrata* and to a lesser extent to *C. albicans*. In ROS induction, killing assays, and western blot analyses of Pyk2 phosphorylation, *C. glabrata* was consistently more dependent on SKAP2 than was *C. albicans* (Figures 5–6). These differences could be attributed to different morphologies and/or cell wall composition (Brunke and Hube, 2013, Dujon et al., 2004). Prior work using human neutrophils showed that *C. albicans* and *C. glabrata* induce different levels of neutrophil activation, ROS, and phagocytosis (Duggan et al., 2015), potentially due to differences in the N-mannan structures, and the glucan exposure geometries at the molecular level (Graus et al., 2018). This is consistent with the idea these pathogens do not trigger identical receptors on neutrophils and that signaling from some of these receptors is more dependent on SKAP2 than from others. Nonetheless, work using knockout mice showed that both Dectin-1 and Dectin-2 greatly contribute to the protection of mice against systemic *C. albicans* and *C. glabrata* infection (Thompson et al., 2019, Ifrim et al., 2014). Thus, our observations that SKAP2 is important for ROS production and adhesion after stimulation by Dectin-1 and Dectin-2 ligands (Figures 2–3, 5) suggests that *Skap2*−/− deficiencies may lead to attenuated responses against systemic *C. albicans* and *C. glabrata*.

Prior work and our data here revealed that infection of neutrophils with *C. glabrata, C. albicans* or *K. pneumoniae* triggers signaling cascade(s) that requires SFKs, Btk, SKAP2, and that SKAP2-mediates pathway(s) leading to ROS production (Figure 5–6) (Nguyen et al., 2020, Gazendam et al., 2014). However, after infection with these pathogens or stimulation with CLRs agonist, we detected distinct differences in the requirements for SKAP2 for phosphorylation of Syk and Pyk2 (Figure 6G-R, Figure 7, Figure S3 and (Nguyen et al., 2020)). Previous work had shown that Syk tyrosine phosphorylation is required signaling downstream of many receptors including CLRs and integrin, while Pyk2 is required for integrin-mediated degranulation, but not ROS production (Kamen et al., 2011, Zarbock et al., 2007, Mocsai et al., 2002, Mocsai et al., 2003, Gazendam et al., 2014, Futosi, 2013). Since SKAP2 was not required for Syk phosphorylation after CLR, integrin, *C. albicans* or *C. glabrata* stimulation, but was required after *K. pneumoniae* stimulation (Figure 6, Figure S6C, and (Shaban et al., 2020, Nguyen et al., 2020)), the receptor(s) that predominantly sense *K. pneumoniae* infection requires SKAP2 prior to Syk and Pyk2 phosphorylation (Figure 7) and are distinct from those that recognize *C. glabrata* and *C. albicans*. Such receptors could include G-protein-coupled receptors (GPCRs), which have been shown to require SKAP2 for ROS and other functions, but are unlikely to be integrin or the CLRs studied here (Boras et al., 2017, Sharma et al., 2017, Sharma et al., 2014, Castillo et al., 2019, Shaban et al., 2020). By contrast, the receptors that recognize *C. glabrata* and *C. albicans* as well as CLRs trigger activation Syk prior to, or potentially independently, of SKAP2 activation (Figure 7). Since SKAP2 was required for maximal phosphorylation of Pyk2 after *C. glabrata* infection but not after stimulation of Mincle or Dectin-2 (Figure 6M-R), at least one other *C. glabrata* receptor requires SKAP2 for Pyk2 phosphorylation and this potentially contributes to the stronger SKAP2-dependent fungicidal activities observed for *C. glabrata* (Figure 5D-E).

**Figure 7:**
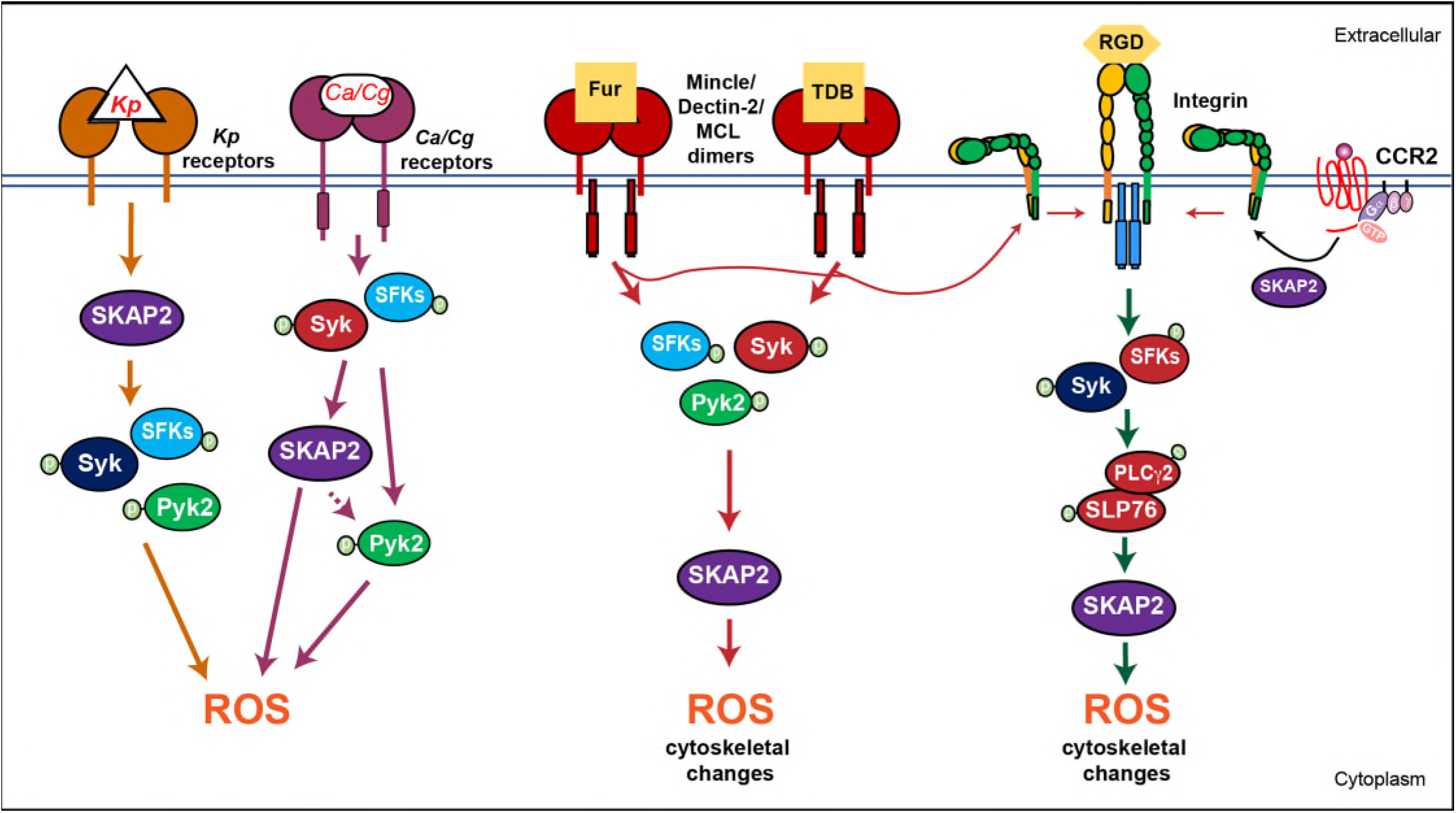
SKAP2 functions prior to or after Syk activation after infection with different pathogens. The binding of *K. pneumoniae* to surface receptors on neutrophils leads to the SKAP2-mediated phosphorylation of Syk and/or Pyk2 and requires the production of extracellular ROS. The binding of *C. glabrata* and *C. albicans* does not require SKAP2 for Syk phosphorylation; however, Pyk2 phosphorylation was dependent on SKAP2. Binding by the CLR agonist, furfurman and TBD induces Syk and Pyk phosphorylation independently of SKAP2. Likewise SKAP2 contributes to extracellular ROS production and cell adhesion following the stimulation of integrins and fMLP (Nguyen et al., 2020, Boras et al., 2017, Shaban et al., 2020).

In summary, our findings broaden the spectrum of PRRs that depend on SKAP2 for activation of neutrophil ROS production and extend the role of SKAP2 in host defense beyond antibacterial immunity to include *Candida* species. Elucidating the mechanism(s) by which SKAP2 differently mediates signaling circuits leading to the activation of neutrophil ROS production after exposure to diverse pathogens is a focus of our future studies. There are significant differences between murine and human neutrophils including the expression of some receptors (Sun et al., 2016, Hajjar et al., 2010, Bagaitkar et al., 2012, Bengoechea and Sa Pessoa, 2019). However, the crucial function of SKAP2-mediated ROS in murine neutrophils following CLRs, *C. albicans*, *C. glabrata,* and *K. pneumoniae* stimulation, and the findings that ROS requires Syk, SFK and Btk in both human and murine neutrophils suggests that parallels exist in signaling through CLRs (Figure 6–7). Therefore, these responses may also depend on SKAP2 in humans. Further linkage studies showing association of Skap2 with autoimmune diseases (Barrett et al., 2009, Jostins et al., 2012), with over 600 *Skap2* variants/mutations reported in the human population across many age and ethnic groups (Karczewski et al., 2020), indicates that analysis of SKAP2 in human neutrophils and other cells is warranted.

## Materials and Methods

### Animals

BALB/c and C57BL/6J mice were purchased from Taconic Biosciences, and Jackson laboratory (Bar Harbor, ME), respectively. Generation of *Skap2-/-* mice in the BALB/c background was previously described (Togni et al., 2005, Alenghat et al., 2012) and were a gift of Dr. Kenneth Swanson. All mice were handled in accordance with protocols approved by the Institutional Animal Care and Use Committee of Tufts University.

### Bacterial and fungal strains

Wild type *C. albicans* (SC5314) and *C. glabrata* (ATCC 2001) were obtained from Michael Mansour lab (MGH, Boston) (Negoro et al., 2020, Xu et al., 2018). Yeast cultures were grown overnight in liquid YPD, and the inoculum prepared as previously described (Negoro et al., 2020, Xu et al., 2018). MKP220, a streptomycin-resistant, capsule type K2 derivative of *K. pneumoniae* ATCC43816 was grown overnight in liquid LB at 37°C with aeration, and the inoculum were prepared as previously described (Silver et al., 2019, Nguyen et al., 2020, Paczosa et al., 2020).

### Neutrophil transfer experiment

Generation of DIV neutrophils and depletion of neutrophils via intraperitoneal injection with α-Ly6G antibody (clone: 1A8) 16 hours prior to infection was completed as previously described (Nguyen et al., 2020) with the following modifications. Hoxb8-immortalized granulocyte myeloid progenitor cells (GMP) were immortalized from C57BL/6J wild-type or *Skap2-/-* mice. Aliquots of ER-HoxB8 GMP were washed 3 times in PBS and cultured in estrogen-free complete RPMI, cRPMI (RPMI-1640 with 10% FBS, 2 mM L-glutamine, and 100U penicillin and 0.1 mg/ml streptomycin) supplemented with 10ng/ml SCF, IL-3, and GCSF for four days. On day 4, DIV neutrophils were collected, pelleted at 250 *x g* for 5 min at 4°C, washed twice with cold sterile PBS, and resuspended in cold sterile PBS at a concentration of 5 × 10^7^ cells/ml. 150ul of cells or sterile PBS were intravenously injected into neutropenic mice. An hour later, mice were intranasally infected with 5000 CFU of *K. pneumoniae*. Mice were sacrificed after 24 hours and lungs harvested for bacterial burden as described (Nguyen et al., 2020).

### Analysis of C-type lectin receptors by flow cytometry

BALB/c wild-type or *Skap2-/-* were infected with *K. pneumoniae*, and lungs were harvested at 24 hours post-infection and were processed as previously described (Nguyen et al., 2020). Dectin-1, Mincle, and Dectin-2 expression were examined by using rat anti-mouse Dectin-1 (Biolegend) followed by Alexa-594-conjugated goat anti-rat (Invitrogen), rabbit anti-mouse Mincle (MyBioSource) followed by Alexa-594-labeled goat anti-rabbit (Invitrogen), or biotinylated goat-anti-mouse Dectin-2 (R&D Systems) followed by APC-conjugated streptavidin antibodies (Biolegend). FlowJo software (Tree Star) was used to analyze all data.

### Neutrophil ROS assays

The isolation of bone marrow neutrophils from BALB/c wild-type (WT) or *Skap2-/-* mice, generation of Hoxb8-immortalized granulocyte myeloid progenitor cells, differentiation of the DIV neutrophils, and ROS assays were done as previously described (Nguyen et al., 2020) with the following modifications. For C-type lectin (CLR) stimulation, 4HBX-96 well plates (Fisher Scientific) were coated with 25μg/ml TDB (InvivoGen), 10μg/ml furfurman (InvivoGen), or 100μg/ml curdlan (InvivoGen) for 3 hours at 37°C, and washed once with PBS before used. For controls, cells were stimulated with 100nM of PMA (Nguyen et al., 2020). Superoxide detection by cytochrome C (Sigma) was performed as previously described (Nguyen et al., 2020, Shaban et al., 2020, Dahlgren et al., 2020). For inhibitor studies, DIV neutrophils were pre-treated with 10nM PP2 (Selleck Chemical), 2μM R406 (Selleck Chemical), 1μM Ibrutinib (Selleck Chemical) for 10 minutes, or DMSO (1:10,000 final concentration, Sigma) as the vehicle control as previously described (Nguyen et al., 2020). To detect ROS production following *Candida* exposure, an isoluminol assay was performed as previously described (Kobayashi et al., 2016, Nguyen et al., 2020, Dahlgren et al., 2020) with the following modifications. *C. albicans* and *C. glabrata* were added to wells containing neutrophils at an MOI of 1, and the plate was centrifuged at 500 *x g* for 3 minutes at 4°C. A BioTek Synergy HT late reader was used to detect absorbances at 490nm and 550nm for cytochrome C assays, or chemiluminescence (RLU) for isoluminol assays. RLU values shown are after subtracting the uninfected samples. Total ROS production was calculated by the sum of the area under the curves for the indicated time in figure legends.

### Spreading assay

DIV neutrophils (1 × 10^5^ cells/well) in HBSS with Ca^2+^ and Mg^2+^ (HBSS+) were plated onto 96-well 4HBX plates coated with 10% FBS, 25μg/ml TDB, 10μg/ml furfurman, 100μg/ml curdlan, or immobilized IgG immune complexes (IC, (Nguyen et al., 2020)) in technical duplicates, and the plate was spun at 250 *x g* for 2 minutes. Cells were incubated at 37°C in presence of 5% CO_2_ for 1 hour. The cells were washed twice with room temperature (RT) PBS, and fixed with 4% formaldehyde for 15 minutes at RT, washed with 200μl of sterile PBS, and then the plate was covered with parafilm and stored at 4°C. The cells were imaged using a Keyence digital bright-field microscope at 20X magnification, and the images were blinded, and assessed for the number of non-spread and spread cells. Three random images were taken, and at least 200 cells were counted per well. The percent of adhesion was calculated as follows: (number of spreading cells)/total number of cells counted for each replicate; an average of technical replicates was calculated for each experiment, and normalized to the unstimulated level. The average of TDB, furfurman, or curdlan-stimulated total adhesion was then divided by the average IC-stimulated adhesion within the respective experiment. Data shown is compiled from 3-4 independent experiments.

### Integrin activation

DIV neutrophils (1 × 10^5^ cells/well) in HBSS+ were plated onto 96-well 4HBX plates coated with 10% FBS, 25μg/ml TDB, 10μg/ml furfurman, or 100μg/ml curdlan and the plate was spun at 250 *x g* for 2 minutes. Negative controls were treated with 2.5mM EDTA for 30 minutes at 37°C and plated in wells coated with 10% FBS. Cells were incubated at 37°C in the presence of 5% CO_2_ for 20 minutes. 30 μg/ml Alexa 488-conjugated fibrinogen (Invitrogen) was added to the wells, and the cells were incubated at 37°C in presence of 5% CO_2_ for an additional 20 minutes. In some FBS-coated wells, 2.5mM manganese (Mn^2+^) was added to cells for the last 15 minutes of incubation. Following stimulation, the supernatants were removed, and the wells were washed twice with cold PBS. 200μl of cold 2.5 mM EDTA in PBS without Ca^2+^ and Mg^2+^ was then added to each well. The cells were incubated at 4°C for an hour to promote lifting of the cells and transferred into 5 ml FACS tubes with 1% formaldehyde. The FACS tubes were covered with paraffin, and stored in the dark, at 4°C until flow cytometry analysis. The percentage of fibrinogen positive cells were normalized to EDTA-treated level and presented as a percentage of Mn^2+^-treated level.

### Candida viability assay

The PrestoBlue assay kit (Thermo Fisher) was used to measure fungal viability following co-culture with WT and *Skap2-/-* DIV neutrophils (Tam et al., 2019, Xu et al., 2018, Negoro et al., 2020). DIV neutrophils in cRPMI were plated in triplicate 96-well non-tissue culture treated plates at a density of 1 × 10^5^ cells/well. WT and *Skap2-/-* DIV neutrophils were stimulated with *C. albicans* or *C. glabrata* (MOI of 0.2), and the plate was spun at 500 *x g* for 30 seconds at RT. The plate was incubated at 37°C in the presence of 5% CO_2_ for 2 or 18 hours for *C. albicans* and *C. glabrata*, respectively. At the end of the incubation period, the plate was centrifuged at 500 *x g* for 3 minutes at 4°C. The supernatant was removed, and the cells were lysed with NP-40 for 5 minutes at 4°C. The lysates were incubated with PrestoBlue Cell Viability Reagent with fresh cRPMI for the *C. albicans,* or YPD broth *C. glabrata*. Fluorescence readings at 560/590nm using a SpectraMax i3x reader (Molecular Devices) or BioTek Synergy H1 at 30°C for at least 18 hours were recorded. PrestoBlue outgrowth data was assessed using a nonlinear four-parameter curve fit equation in GraphPad Prism 7 to extract the 50% maximal value (MIC_50_) for the calculation of the number of yeasts in each well (Xu et al., 2018, Negoro et al., 2020). Controls included in each plate/experiment were media only wells and a standard of *C. albicans* or *C. glabrata* (no neutrophils), which was prepared by serially diluting the yeasts starting at MOI 2. Data from the *Candida-*only wells were fitted to linear regression curve, and used to calculate the number of surviving yeasts that were co-incubated with or without neutrophils. Percent of yeast killing was calculated as follows: 100-(100*viability of yeast plus neutrophils /viability of yeast only).

### Western blot analysis

Western blot analysis was conducted as previously described (Nguyen et al., 2020, Shaban et al., 2020) with the following modifications. For purified agonists, DIV neutrophils were plated onto 96-well 4HBX plates coated with IC, 25 μg/ml TDB, 10 μg/ml furfurman, or 250 μg/ml curdlan. DIV neutrophils (7.5×10^5^ cells/well) in HBSS+ were plated onto 96-well 4HBX plates coated with 10% FBS. *C. albicans* and *C. glabrata* were added at MOI 2, while *K. pneumoniae* was added at a MOI of 40; the plate was spun at 250 *x g* for 2 minutes. The cells were incubated at 37°C in the presence of 5% CO_2_ for 15 minutes, lysed in 1X Novex buffer and 5-7.5×10^5^ cells equivalents were resolved on 4-12% NuPAGE gel (Invitrogen) in MOPS buffer. Antibodies to phosphorylated proteins include Syk-Y352 (Cell Signaling Technology), and Pyk2-Y402 (Cell Signaling Technology). All blots were also probed with RhoGDI (Cell Signaling Technology). Secondary LI-COR goat anti-mouse or goat anti-rabbit IRDye 800 CW (Cell Signaling Technology) were used at a dilution of 1:20,000. The blots were then stripped as previously described (Nguyen et al., 2020, Shaban et al., 2020), and re-probed with antibodies for total proteins against Syk, and Pyk2 (Cell Signaling Technology). The Odyssey CLx LI-COR system and IS Image Studio Lite were used for analysis of the blots and quantification of bands. The normalized protein levels of pSyk, and pPyk2 were calculated by taking the ratio of phosphorylated to total protein and then normalizing to the RhoGDI loading control. Normalized phospho-protein levels were used to calculate the fold change to unstimulated control within each group, or to unstimulated control of WT (Supplementary file). In addition, the relative induction of phosphorylation in the unstimulated controls was determined by dividing the fold change of each experiment by the average of the three experiments. Fold change was log transformed and subjected to statistical analysis.

### Statistics

All statistical analyses were performed using Graph Pad Prism version 7.

## Acknowledgements

This work was supported by NIH NIAID R01 AI113166 awarded to JM and NIH NIAID RO1 AI132638 to MKM. The authors declare no competing financial interests. We thank Lamyaa Shaban, Alyssa Fasciano, Rebecca Silver, Anne McCabe, Michelle Sodipo, Parisa Kalantari, Miles Duncan, and Maria-Cristina Seminario for critically reading the manuscript and/or for helpful scientific and technical discussions.

## Supplemental Figures and Tables

**Supplemental figure 1. Neutrophils from *K. pneumoniae*-infected lungs are primarily Mincle-positive, Dectin-1hi, and Dectin-2hi cells.** BALB/c wild-type or *Skap2-/-* mice were infected intranasally with *K. pneumoniae,* and lungs were harvested after 24 hours. (**A-C, E-H**) Analysis of CLR expression on neutrophils (CD11b+ Ly6G^hi^). Cells were stained with CD11b and Ly6G, fixed with 4% formaldehyde, and stained extracellularly or intracellularly with α-Mincle, α-Dectin-2, or α-Dectin1 as described in the *Materials and Methods*. Neutrophils (CD11b^+^ Ly6G^hi^) were gated to analyze for expression of Mincle, Dectin-2, and Dectin-1. **(A-C)** Representative dot plots from a *K. pneumoniae*-infected WT mouse. **(D)** CFU of *K. pneumoniae* recovered from infected mice. Each dot represents a mouse, and black bar represents geometric mean. **(E-H)** Dotted line indicate average level in PBS-treated lungs. Data are compiled from **(F)** 1 experiment, or **(D-E, G-H)** 2 independent experiments with 3 mice/group. **(D-H)** Significance was assessed using two-tail unpaired Student’s *t* test; log-transformed numbers were used for **(D)**.

**Supplemental Figure 2: BM neutrophils require SKAP2 for TDB, furfurman, and curdlan-induced ROS.** The respiratory burst of BALB/c (WT) and *Skap2-/-* DIV **(A-C)** or BM **(D-F)** neutrophils plated on immobilized TDB, furfurman (Fur), or curdlan (Cur). **(G-H)** The total concentration of superoxide produced by BM neutrophils after 60 minutes was calculated by the sum of the area under the curves, and presented as **(G)** total superoxide (normalized to unstimulated values), or **(H)** as percentage of PMA-induced ROS. **(A-C)** A representative experiment performed in triplicate of superoxide production from n=8-9 and presented as mean + SD. **(D-H)** Data are presented as mean + SD of technical triplicates from n=1. Statistics were analyzed by **(A-F)** two-way ANOVA with Tukey’s post-test, or **(G-H)** one-way ANOVA with Sidak’s.

**Supplemental figure 3: SKAP2 is required for maximal *C. albicans* and *C. glabrata*-induced phosphorylation of Pyk2.** WT and *Skap2-/-* DIV neutrophils were stimulated with MOI 2 of *C. albicans* and *C. glabrata* for 15 min. **(A-B)** Induced phosphorylation levels of Syk, and Pyk2 are shown as mean + SEM with each experiment represented by a different color. Solid symbols indicate the representative blots shown in Figure 6. Significance was assessed using one-way ANOVA with Sidak’s. **(C)** Western Blot of WT and *Skap2-/-* neutrophils were plated onto immobilized TDB, or IgG immune complexes (IC) or infected with K. pneumoniae (MOI of 40) for 15 minutes at 37°C. Lysates were analyzed by western blot for pSyk (Y352), pPyk2 (Y402), and RhoGDI. Blots were then stripped and re-probed for total Syk, and Pyk2. Phospho-signal was first divided by total protein, then by loading control, and then normalized to unstimulated values as described in the *Materials and Methods*.

