## Supplemental Figures and Tables for "Neutrophils require SKAP2 for reactive oxygen species production following C-type lectin and *Candida* stimulation"

### Figure 1 – supplemental figure 1

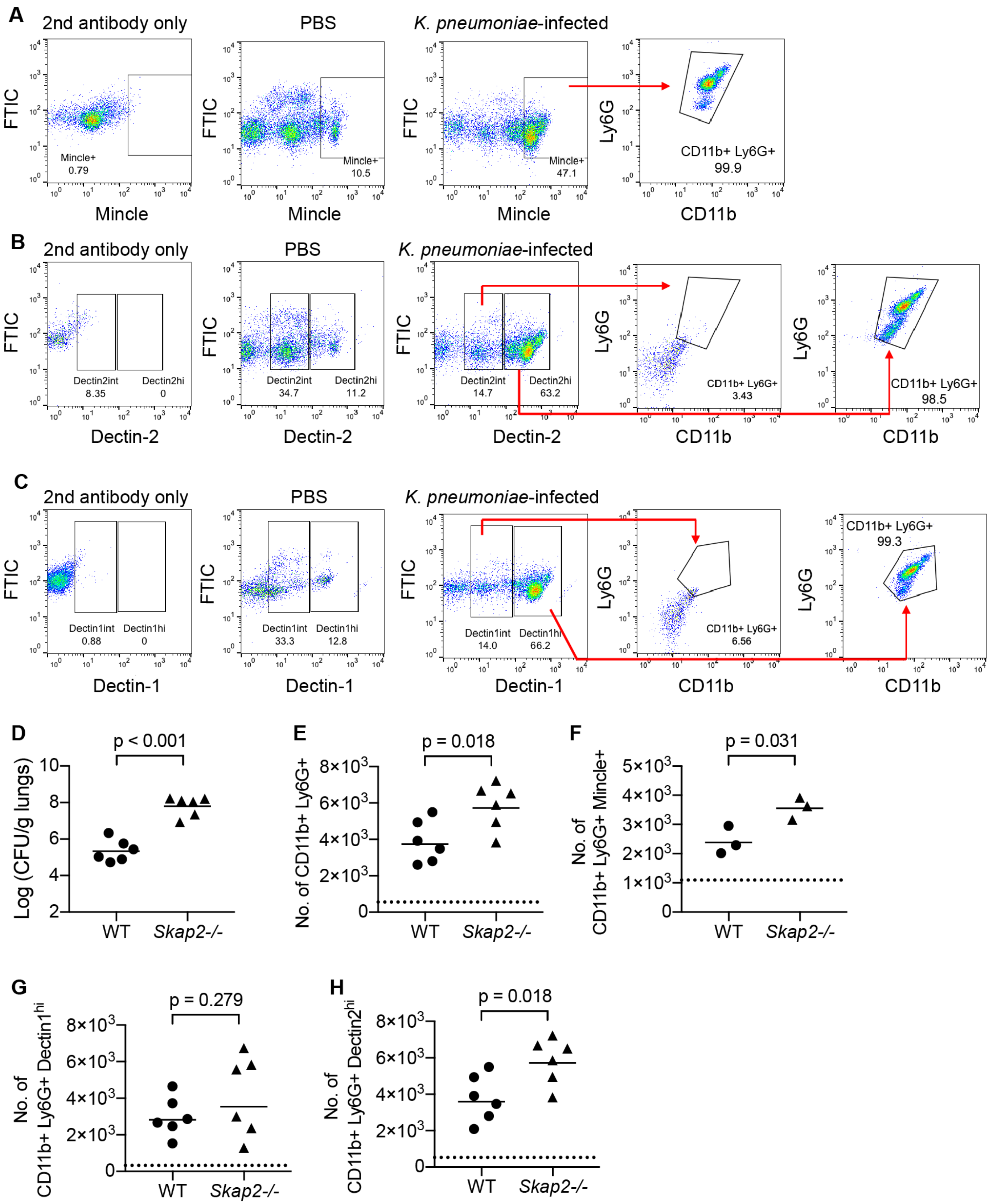

**Figure 2 – supplemental figure 1**

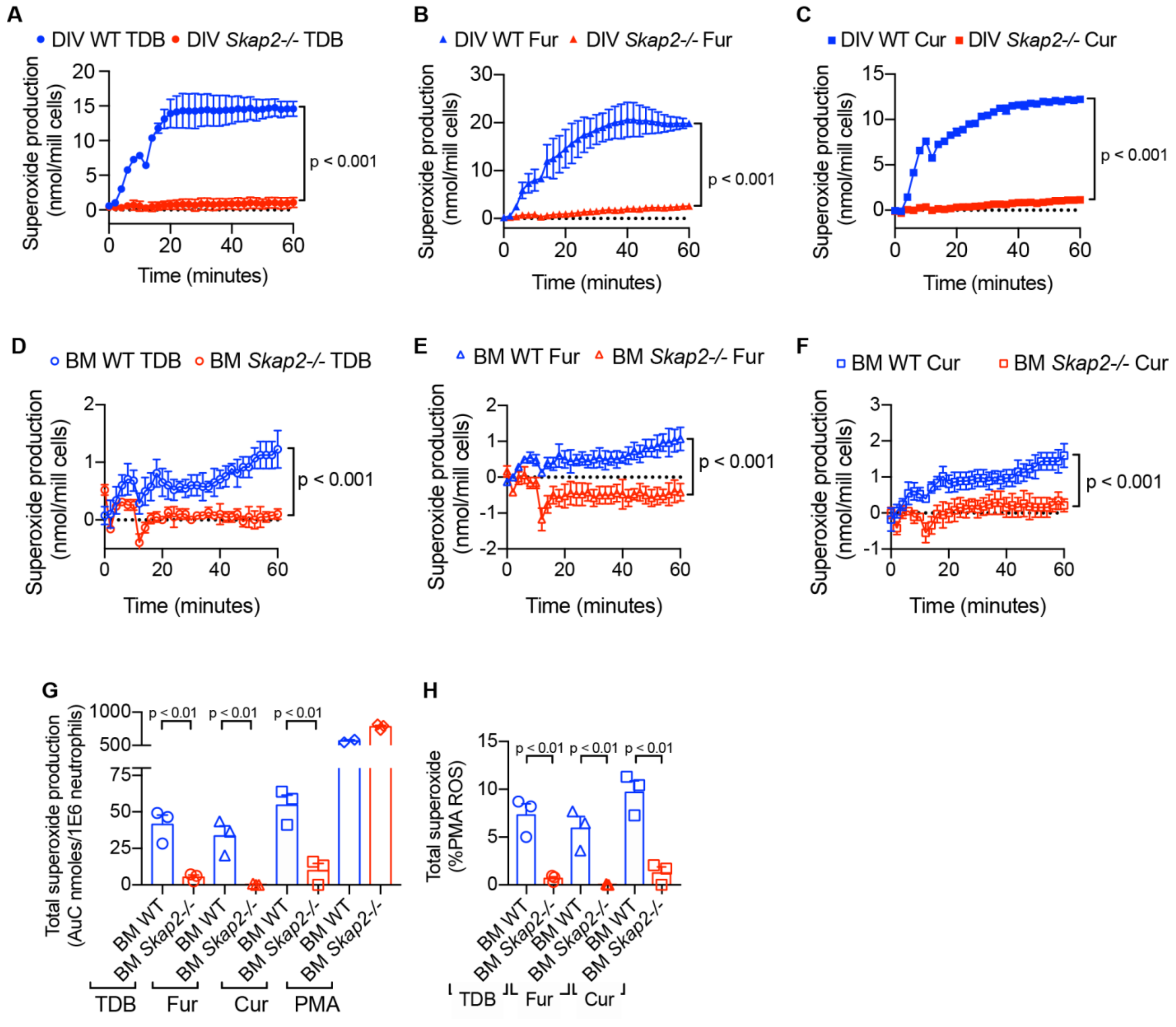

**Figure 6 – supplemental figure 1**

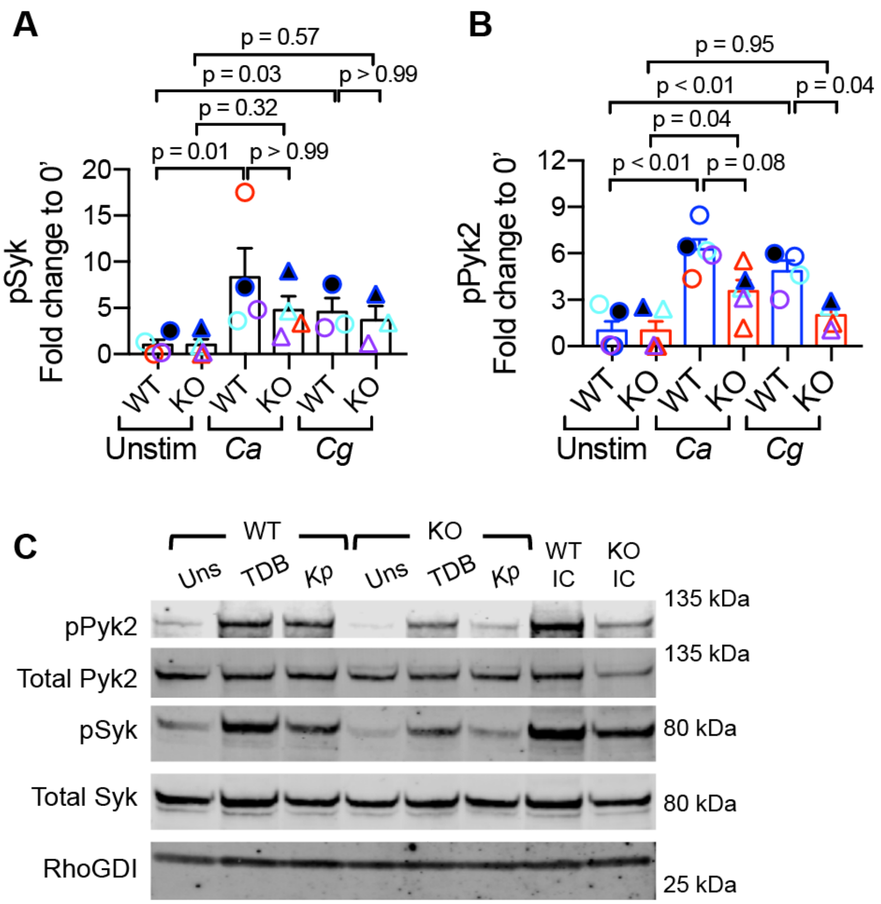

Table S1: Calculations for the induction of TDB and furfurman (Fur)-induced phospho-Syk.

| | Normalized phospho-Syk intensity[b] | | | | | | | |
|---|---|---|---|---|---|---|---|---|
| | WT | | | | *Skap2-/-* | | | |
| | Uns | TDB | Fur | IC | Uns | TDB | Fur | IC |
| Experiment 1 | 3.72 | 6.63 | | 13.87 | 4.53 | 8.57 | | 18.26 |
| Experiment 2 | 5.04 | 13.23 | 9.07 | 21.85 | 1.35 | 6.03 | 8.23 | 21.85 |
| Experiment 3 | 1.55 | 8.16 | 9.26 | 13.91 | 1.20 | 4.81 | 4.50 | 11.94 |
| Experiment 4 | 0.16 | 1.16 | 1.20 | 1.31 | 0.20 | 1.42 | 1.62 | 2.28 |
| | Fold change of phospho-Syk to uns within each group[b] | | | | | | | |
| | WT | | | | *Skap2-/-* | | | |
| | Uns | TDB | Fur | IC | Uns | TDB | Fur | IC |
| Experiment 1 | 1.00 (1.42) | 1.78 | | 3.73 | 1.00 (2.49) | 1.89 | | 4.03 |
| Experiment 2 | 1.00 (1.93) | 2.62 | 1.80 | 4.33 | 1.00 (0.74) | 4.47 | 6.11 | 16.21 |
| Experiment 3 | 1.00 (0.59) | 5.28 | 5.99 | 9.01 | 1.00 (0.66) | 4.02 | 3.76 | 9.99 |
| Experiment 4 | 1.00 (0.06) | 7.43 | 7.69 | 8.40 | 1.00 (0.11) | 7.11 | 8.11 | 11.41 |
| | Fold change of phospho-Syk to WT Uns[d] | | | | | | | |
| | WT | | | | *Skap2-/-* | | | |
| | Uns | TDB | Fur | IC | Uns | TDB | Fur | IC |
| Experiment 1 | 1.00 | 1.78 | | 3.73 | 1.22 | 2.31 | | 4.91 |
| Experiment 2 | 1.00 | 2.62 | 1.80 | 4.33 | 0.27 | 1.19 | 1.63 | 4.33 |
| Experiment 3 | 1.00 | 5.28 | 5.99 | 9.01 | 0.77 | 3.11 | 2.91 | 7.73 |
| Experiment 4 | 1.00 | 7.43 | 7.69 | 8.40 | 1.28 | 9.11 | 10.40 | 14.63 |

WT and *Skap2-/-* DIV neutrophils were stimulated with TDB, furfurman (Fur), or IgG IC (as positive control) for 10 minutes at 37ºC. Lysates were prepared and analyzed by western blot for phospho-Syk, and RhoGDI (loading control). Blots were then stripped and re-probed for the respective total protein. Quantification of protein level was assessed using Licor.

a Normalized phospho-Syk = ratio of the intensity of phosphorylated bands to total protein to RhoGDI loading control.

b Fold change was calculated by diving the normalized values of stimulated samples by the normalized value of unstimulated (uns) sample within respective group of each experiment.

c Fold change in parenthesis was calculated by diving the normalized values of unstimulated samples of each experiment by the average normalized value of unstimulated (uns) sample from all three experiments.

d Fold change was calculated by diving the normalized values of stimulated samples by the normalized value of WT unstimulated (uns) sample of each experiment.

Table S2. Calculations for the induction of *C. albicans* and *C. glabrata*-induced phospho-Syk.

| | Normalized phospho-Syk intensity[b] | | | | | | | |
|---|---|---|---|---|---|---|---|---|
| | WT | | | | *Skap2-/-* | | | |
| | Uns | *Ca* | *Cg* | IC | Uns | *Ca* | *Cg* | IC |
| Experiment 1 | 2.13 | 7.66 | 7.00 | 14.48 | 1.53 | 7.12 | 5.19 | 15.14 |
| Experiment 2 | 4.08 | 29.66 | 30.79 | 13.36 | 4.60 | 41.49 | 29.97 | 9.47 |
| Experiment 3 | 0.01 | 0.22 | | 0.09 | 0.01 | 0.03 | | 0.13 |
| Experiment 4 | 0.27 | 1.30 | 0.77 | 2.02 | 0.32 | 0.62 | 0.37 | 4.52 |
| | Fold change of phospho-Syk to uns within each group[b] | | | | | | | |
| | WT | | | | *Skap2-/-* | | | |
| | Uns | *Ca* | *Cg* | IC | Uns | *Ca* | *Cg* | IC |
| Experiment 1 | 1.00 (1.31) | 3.60 | 3.29 | 6.81 | 1.00 (0.95) | 4.66 | 3.40 | 9.91 |
| Experiment 2 | 1.00 (2.51) | 7.28 | 7.55 | 3.28 | 1.00 (2.85) | 9.02 | 6.52 | 2.06 |
| Experiment 3 | 1.00 (0.01) | 17.52 | | 7.42 | 1.00 (0.001) | 3.39 | | 16.01 |
| Experiment 4 | 1.00 (0.17) | 4.81 | 2.85 | 7.45 | 1.00 (0.20) | 1.93 | 1.16 | 14.05 |
| | Fold change of phospho-Syk to WT Uns[d] | | | | | | | |
| | WT | | | | *Skap2-/-* | | | |
| | Uns | *Ca* | *Cg* | IC | Uns | *Ca* | *Cg* | IC |
| Experiment 1 | 1.00 | 3.60 | 3.29 | 6.81 | 0.72 | 3.35 | 2.44 | 7.12 |
| Experiment 2 | 1.00 | 7.28 | 7.55 | 3.28 | 1.13 | 10.18 | 7.35 | 2.32 |
| Experiment 3 | 1.00 | 17.52 | | 7.42 | 0.62 | 2.11 | | 9.97 |
| Experiment 4 | 1.00 | 4.81 | 2.85 | 7.45 | 1.19 | 2.29 | 1.38 | 16.69 |

WT and *Skap2-/-* DIV neutrophils were infected with *C. albicans* , *C. glabrata* or stimulated IgG IC (as positive control) for 15 minutes at 37°C. Lysates were prepared and analyzed by western blot for phospho-Syk, and RhoGDI (loading control). Blots were then stripped and re-probed for the respective total protein. Quantification of protein level was assessed using Licor.
a Normalized phospho-Syk = ratio of the intensity of phosphorylated bands to total protein to RhoGDI loading control.
b Fold change was calculated by diving the normalized values of stimulated samples by the normalized value of unstimulated (uns) sample within respective group of each experiment.
c Fold change in parenthesis was calculated by diving the normalized values of unstimulated samples of each experiment by the average normalized value of unstimulated (uns) sample from both experiments.
d Fold change was calculated by diving the normalized values of stimulated samples by the normalized value of WT unstimulated (uns) sample of each experiment.

Table S3: Calculations for the induction of TDB and furfurman (Fur)-induced phospho-Pyk2.

| | Normalized phospho-Pyk2 intensity[a] | | | | | | | |
|---|---|---|---|---|---|---|---|---|
| | WT | | | | ***Skap2-/-*** | | | |
| | **Uns** | **TDB** | **Fur** | **IC** | **Uns** | **TDB** | **Fur** | **IC** |
| Experiment 1 | 25.59 | 48.65 | | 74.06 | 25.69 | 84.01 | | 334.16 |
| Experiment 2 | 18.66 | 86.46 | 95.48 | 143.07 | 19.52 | 70.28 | 74.48 | 107.69 |
| Experiment 3 | 40.49 | 282.33 | 200.87 | 404.76 | 45.27 | 276.67 | 128.58 | 433.30 |
| Experiment 4 | 0.23 | 1.91 | 1.92 | 1.95 | 0.32 | 1.75 | 2.18 | 3.45 |
| | Fold change of phospho-Pyk2 to uns within each group[b] | | | | | | | |
| | **WT** | | | | ***Skap2-/-*** | | | |
| | **Uns** | **TDB** | **Fur** | **IC** | **Uns** | **TDB** | **Fur** | **IC** |
| Experiment 1 | 1.00 (1.21) | 1.90 | | 2.89 | 1.00 (1.13) | 3.27 | | 13.01 |
| Experiment 2 | 1.00 (0.88) | 4.63 | 5.12 | 7.67 | 1.00 (0.86) | 3.60 | 3.82 | 5.52 |
| Experiment 3 | 1.00 (1.91) | 6.97 | 4.96 | 10.00 | 1.00 (1.99) | 6.11 | 2.84 | 9.57 |
| Experiment 4 | 1.00 (0.01) | 8.32 | 8.36 | 8.50 | 1.00 (0.01) | 5.45 | 6.80 | 10.76 |
| | Fold change of phospho-Pyk2 to WT Uns[d] | | | | | | | |
| | **WT** | | | | ***Skap2-/-*** | | | |
| | **Uns** | **TDB** | **Fur** | **IC** | **Uns** | **TDB** | **Fur** | **IC** |
| Experiment 1 | 1.00 | 1.90 | | 2.89 | 1.00 | 3.28 | | 13.06 |
| Experiment 2 | 1.00 | 4.63 | 5.12 | 7.67 | 1.05 | 3.77 | 3.99 | 5.77 |
| Experiment 3 | 1.00 | 6.97 | 4.96 | 10.00 | 1.12 | 6.83 | 3.18 | 10.70 |
| Experiment 4 | 1.00 | 8.32 | 8.36 | 8.50 | 1.40 | 7.63 | 9.51 | 15.05 |

WT and *Skap2-/-* DIV neutrophils were stimulated with TDB, furfurman (Fur), or IgG IC (as positive control) for 10 minutes at 37ºC. Lysates were prepared and analyzed by western blot for phospho-Pyk2, and RhoGDI (loading control). Blots were then stripped and re-probed for the respective total protein. Quantification of protein level was assessed using Licor.

a Normalized phospho-Pyk2 = ratio of the intensity of phosphorylated bands to total protein to RhoGDI loading control.

b Fold change was calculated by diving the normalized values of stimulated samples by the normalized value of unstimulated (uns) sample within respective group of each experiment.

c Fold change in parenthesis was calculated by diving the normalized values of unstimulated samples of each experiment by the average normalized value of unstimulated (uns) sample from all three experiments.

d Fold change was calculated by diving the normalized values of stimulated samples by the normalized value of WT unstimulated (uns) sample of each experiment.

Table S4. Calculations for the induction of *C. albicans* and *C. glabrata*-induced phospho-Pyk2

| | Normalized phospho-Pyk2 intensity[a] | | | | | | | |
|---|---|---|---|---|---|---|---|---|
| | **WT** | | | | ***Skap2-/-*** | | | |
| | **Uns** | ***Ca*** | ***Cg*** | **IC** | **Uns** | ***Ca*** | ***Cg*** | **IC** |
| Experiment 1 | 14.47 | 89.09 | 66.39 | 92.90 | 14.83 | 55.53 | 37.75 | 55.63 |
| Experiment 2 | 11.89 | 76.51 | 71.13 | 42.09 | 15.49 | 66.15 | 45.40 | 18.30 |
| Experiment 3 | 0.00 | 0.01 | | 0.00 | 0.00 | 0.00 | | 0.00 |
| Experiment 4 | 0.03 | 0.25 | 0.17 | | 0.03 | 0.18 | 0.05 | |
| Experiment 5 | 0.33 | 1.95 | 0.99 | 1.48 | 0.43 | 1.34 | 0.45 | 4.50 |
| | Fold change of phospho-Pyk2 to uns within each group[b] | | | | | | | |
| | **WT** | | | | ***Skap2-/-*** | | | |
| | **Uns** | ***Ca*** | ***Cg*** | **IC** | **Uns** | ***Ca*** | ***Cg*** | **IC** |
| Experiment 1 | 1.00 (2.71) | 6.16 | 4.59 | 6.42 | 1.00 (2.41) | 3.75 | 2.55 | 3.75 |
| Experiment 2 | 1.00 (2.23) | 6.43 | 5.98 | 3.54 | 1.00 (2.52) | 4.27 | 2.93 | 1.18 |
| Experiment 3 | 1.00 (0.00) | 9.70 | | 2.41 | 1.00 (0.00) | 1.37 | | 6.42 |
| Experiment 4 | 1.00 (0.01) | 8.47 | 5.81 | 0.00 | 1.00 (0.01) | 5.54 | 1.48 | 0.00 |
| Experiment 5 | 1.00 (0.06) | 5.90 | 3.00 | 4.49 | 1.00 (0.07) | 3.09 | 1.05 | 10.42 |
| | Fold change of phospho-Pyk2 to WT Uns[d] | | | | | | | |
| | **WT** | | | | ***Skap2-/-*** | | | |
| | **Uns** | ***Ca*** | ***Cg*** | **IC** | **Uns** | ***Ca*** | ***Cg*** | **IC** |
| Experiment 1 | 1.00 | 6.16 | | 6.42 | 1.02 | 3.84 | | 3.84 |
| Experiment 2 | 1.00 | 6.43 | 5.98 | 3.54 | 1.30 | 5.56 | 3.82 | 1.54 |
| Experiment 3 | 0.00 | 0.00 | 0.00 | 0.00 | 0.00 | 0.00 | 0.00 | 0.00 |
| Experiment 4 | 0.00 | 0.02 | 0.01 | 0.00 | 0.00 | 0.02 | 0.00 | 0.00 |
| Experiment 5 | 0.03 | 0.16 | 0.08 | 0.12 | 0.04 | 0.11 | 0.04 | 0.38 |

WT and *Skap2-/-* DIV neutrophils were infected with *C. albicans*, *C. glabrata* or stimulated IgG IC (as positive control) for 10 minutes at 37°C. Lysates were prepared and analyzed by western blot for phospho-Pyk2, and RhoGDI (loading control). Blots were then stripped and re-probed for the respective total protein. Quantification of protein level was assessed using Licor.

a Normalized phospho-Pyk2 = ratio of the intensity of phosphorylated bands to total protein to RhoGDI loading control.

b Fold change was calculated by diving the normalized values of stimulated samples by the normalized value of unstimulated (uns) sample within respective group of each experiment.

c Fold change in parenthesis was calculated by diving the normalized values of unstimulated samples of each experiment by the average normalized value of unstimulated (uns) sample from both experiments.

d Fold change was calculated by diving the normalized values of stimulated samples by the normalized value of WT unstimulated (uns) sample of each experiment.
